# NMDA receptor-dependent presynaptic homeostatic plasticity?

**DOI:** 10.64898/2026.02.28.708706

**Authors:** Tianli Dou, Junting Zhang, Yidan Hong, X. Chen, R.A. Nicoll

**Author notes:** Address correspondence to: Xiumin Chen, Roger A. Nicoll. These authors contributed equally (X. Chen, R.A. Nicoll).

## Abstract

Excitatory glutamatergic synapses in the brain are remarkably plastic. Two forms of plasticity have received the most attention: long-term potentiation (LTP) and synaptic homeostasis. While LTP requires the activation of NMDA receptors, synaptic homeostasis does not. However, both phenomena are mediated by the recruitment of postsynaptic AMPA receptors to the synapses. Recently a new form of plasticity has been described referred to as presynaptic homeostatic plasticity (PHP) (Chipman et al., 2022; Chipman et al., 2025). Pharmacological inhibition of AMPA synaptic responses in CA1 hippocampal pyramidal cells initiates a rapid homeostatic response that results in the recovery of the AMPA responses to normal values in the continued presence of the inhibitor. Accompanying this recovery is a doubling of the NMDA response which is interpreted as an increase in the release of glutamate. This is provocative since it is the first report claiming that a reduction in AMPA responses triggers an enhancement in NMDA responses. Using three different protocols to monitor synaptic responses we fail to observe any recovery of synaptic responses in the presence of an AMPA inhibitor. Furthermore, there was no enhancement in NMDA responses. Thus, we find no evidence for the presence of PHP at CA1 hippocampal synapses.

## Introduction

Synaptic plasticity in the central nervous system is fundamental to neural development and learning and memory. Glutamatergic excitatory synapses express two prominent forms of synaptic plasticity. Long term potentiation (LTP) induced by brief high frequency stimulation results in a long-lasting NMDA receptor (NMDAR)-dependent enhancement in AMPA receptor (AMPAR) responses (Diering and Huganir, 2018; Malinow and Malenka, 2002; Nicoll, 2017). The second form of plasticity is an NMDAR-independent homeostatic enhancement of AMPAR responses (Ancona Esselmann et al., 2017; Turrigiano, 2008). Both LTP and homeostatic plasticity are mediated by the recruitment of AMPARs to the synapse. Recently, two papers (Chipman *et al*., 2022; Chipman *et al*., 2025) present evidence in CA1 hippocampal pyramidal cells for a new form of plasticity termed presynaptic homeostatic plasticity (PHP). The physiological evidence for PHP is based on two findings. First, in the continuous presence of the AMPAR selective inhibitor GYKI, depressed AMPAR responses return to control levels within an hour (Fig. 1, (Chipman *et al*., 2022)). Second, the NMDAR responses approximately double in size during this time (Fig. 2, (Chipman *et al*., 2022)). It is proposed that inhibition of AMPAR responses triggers a rapid (within minutes), NMDAR-dependent homeostatic response resulting in a retrograde signal causing the presynaptic increase in glutamate release. To our knowledge this is the first report claiming that a reduction in AMPAR responses trigger an increase in NMDAR responses. Numerous studies have reported that inhibiting AMPAR function, either genetically or pharmacologically fail to show any enhancement in NMDAR function (Diaz-Alonso et al., 2017; Lu et al., 2009; Watson et al., 2017). Acute pharmacological inhibition of CaMKII (Tao et al., 2021) reduces AMPAR EPSCs but has no effect on NMDAR EPSCs. Viral expression of PSD-95 RNAi selectively reduces AMPAR function (Ehrlich and Malinow, 2004; Elias et al., 2006; Schluter et al., 2006). TARP deletion (Chen et al., 2000; Hashimoto et al., 1999; Ravi et al., 2022; Rouach et al., 2005) reduces or eliminates AMPAR function with no change in NMDAR function. In the few instances where a change in NMDAR function has been reported it is invariably a reduction. Thus, in contrast to the numerous studies from many labs, it appears that the NMDAR-dependent PHP reported by Chipman et al. (Chipman *et al*., 2022; Chipman *et al*., 2025) is uniquely triggered by GYKI. Given the provocative and unprecedented nature of these findings, we have reexamined the acute actions of GYKI on CA3-CA1 synapses using well established electrophysiological experiments.

**Fig. 1.**
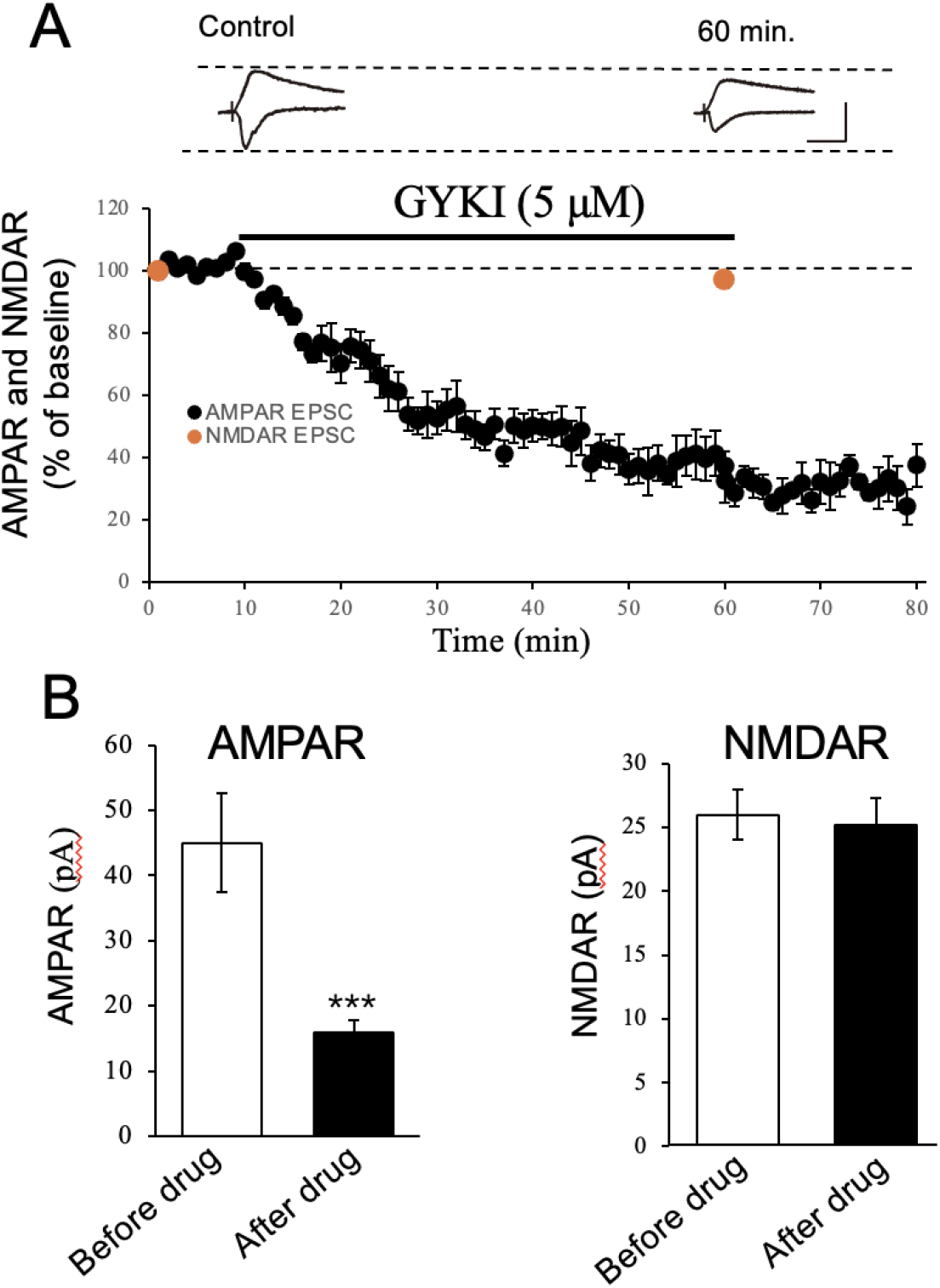
GYKI application causes a persistent depression of AMPAR EPSCs, but no change in NMDAR EPSCs. (A) Sample traces of AMPAR EPSC (inward currents) and NMDAR EPSC (outward currents) before (Control) and after application of GYKI. Graph showing normalized amplitudes of AMPAR EPSCs (black filled circles) and NMDAR EPSCs (red filled circles) before and after GYKI (5 μM) application. n = 6 of cells from 3 mice. (B) Bar graph of AMPAR EPSCs before (44.8 ± 11 pA) and after GYKI (5 μM) (17.5 ± 2 pA, P < 0.001) and NMDAR EPSCs before (26.3 ± 2 pA) and after GYKI (5 μM) (25.2 ± 3 pA). Normalized data were analyzed using a one-way ANOVA followed by the Brown– Forsythe test and Bartlett’s test. (Scale bars, 10 ms, 50 pA.)

**Fig. 2.**
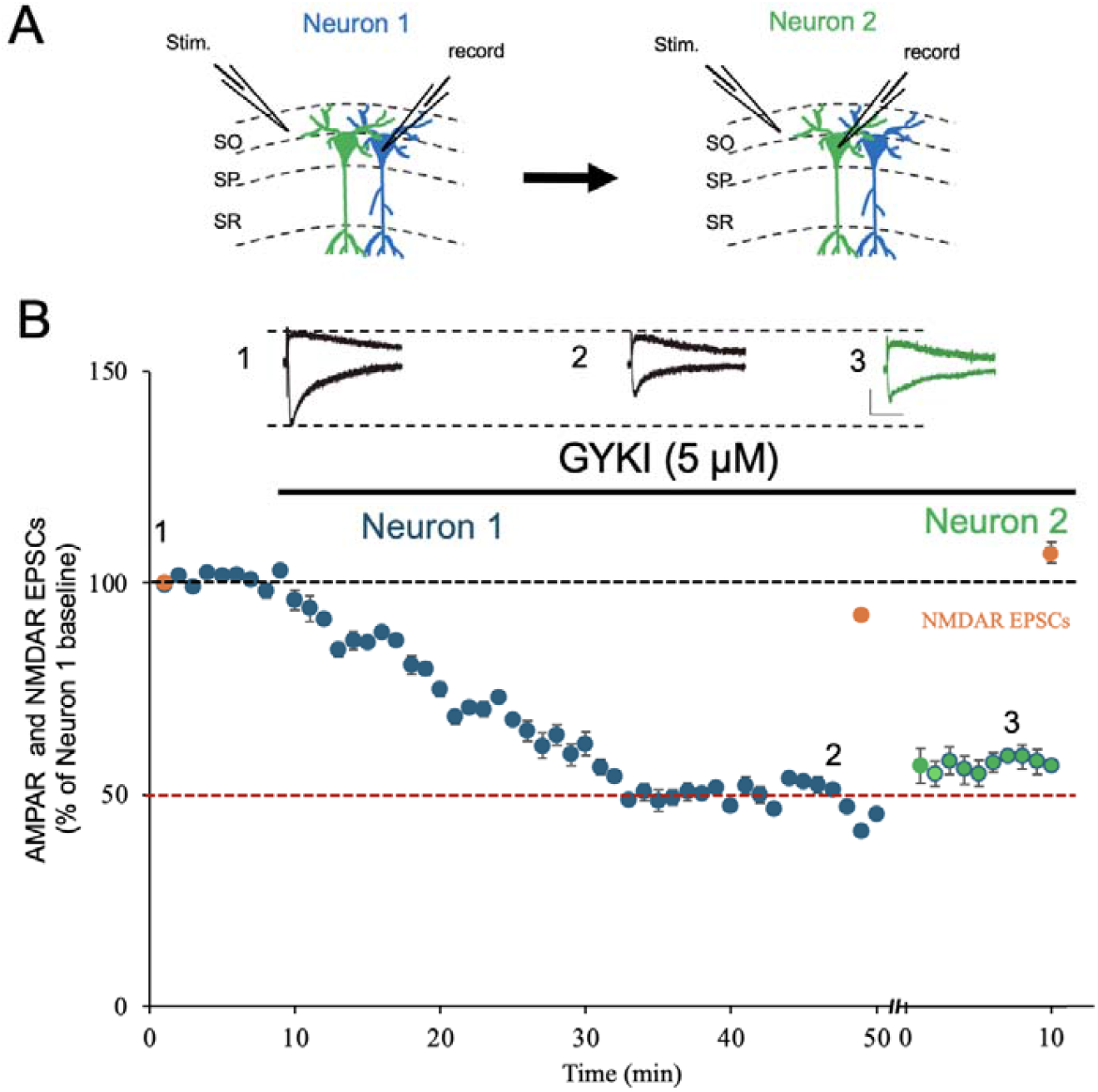
The persistent depression by GYKI is independent of recording conditions. (A) Schematic diagram showing the recording strategy. SO, *stratum oriens*; SP, *stratum pyramidale*; SR, *stratum radiatum*. All experiments are from acute slices. (B) Graph showing normalized amplitudes of AMPAR EPSCs (blue and green filled circles) and NMDAR EPSCs (red filled circles) from two neighboring neurons recorded sequentially. Neuron 1 (blue filled circles) and Neuron 2 (green filled circles) before and after GYKI (5 μM) application. Pairs of two neighboring neurons recorded sequentially (n = 5) from 3 mice. The time between recording from Neuron 1 (blue circles) and Neuron 2 (green circles) varied by a few minutes and time 0 is when recording in Neuron 2 commenced (Scale bars, 10 ms, 50 pA.)

## Results

Whole cell recordings were made from CA1 pyramidal cells in acute hippocampal slices using standard, long established methods (Incontro et al., 2018). A stimulating electrode placed in *Stratum Oriens* was used to activate excitatory synapses. Shortly after establishing the whole cell configuration, the cell was depolarized to +30 mV to record the NMDAR EPSC (measured at 100 ms after the stimulus) (red filled circle in Fig. 1A). The cell was returned to -70 mV and baseline AMPAR responses were recorded (black closed circles in Fig. 1A). GYKI (5 μM) was applied for 50 min. The AMPAR response slowly decreased and stabilized at approximately 40% (Fig. 1A and B). There was no evidence that the action of GYKI diminished during the hour-long period. At this point in the experiment, the cell was depolarized to +30 mV and the NMDAR EPSC recorded. No change in the NMDAR EPSC occurred over the course of the experiment (Fig. 1A and B). Thus, there was no evidence for PHP involving either a recovery of the AMPAR response during the application of GYKI or for a retrograde enhancement of glutamate release, as measured by the NMDAR response. One difference between our experiments and those of Chipman et al. (2022) is that during their experiments they intermittently switched into current clamp. No reason was provided for why this was done. Perhaps this or some other technical issues with whole cell recording during the experiments prevented the PHP in our experiments.

We therefore designed two experiments that circumvent this potential problem. First, we repeated the experiment shown in Fig. 1. However, after recording the responses during a 40 min. exposure to GYKI from one neuron (Fig. 2A, blue neuron), we then record from a neighboring neuron (Fig. 2A, green neuron). If the whole cell recording had interfered with the homeostasis, the AMPAR EPSC in the newly recorded cell should be of the same size as the responses recorded prior to applying GYKI. On the contrary the size of the response in the newly recorded neurons was of the same size as that in the continuously recorded neurons, ruling out an effect of the whole cell recording. The same conclusion can be reached for the NMDAR EPSCs, indicating that there had been no increase in glutamate release.

For the second experiment we turned to traditional field potential recordings. This approach records extracellularly from a large population of neurons and synapses and thus is the least intrusive technique for recording synaptic responses. Furthermore, the recordings are very stable over time. Field potentials were recorded from *Stratum Radiatum* (Fig. 3, inset) or *Stratum Oriens (data not shown)*. Following application of GYKI (5 μM) the AMPAR responses slowly decrease and stabilize after ∼40 min. There was no evidence of any recovery of the responses. These findings complement those obtained in *Stratum Oriens* and extend the findings to synapses in *Stratum Radiatum*, upon which much of our understanding of excitatory synaptic transmission in the CNS is based.

**Fig. 3.**
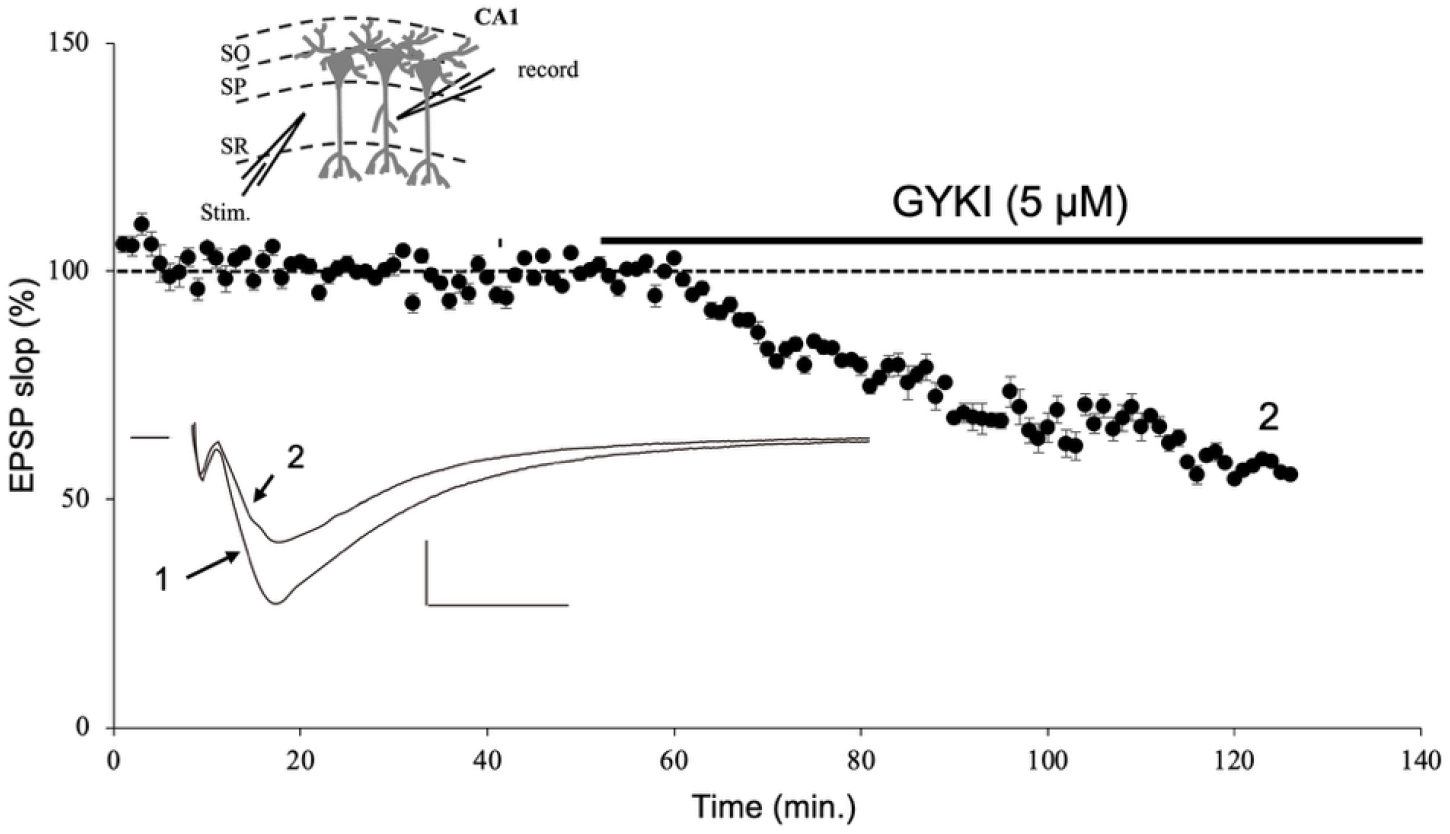
GYKI application causes a persistent depression of field EPSPs. Inset shows schematic diagram of the recording strategy. SO, *stratum oriens*; SP, *stratum pyramidale*; SR, *stratum radiatum*. All experiments are from acute slice. Graph of normalized EPSP slope showing the persistent depression caused by GYKI (5 μM) application. (n = 4 slices from 3 mice) (Scale bars, 10 ms, 0.2 mV) Note that the action of GYKI is slower than the whole cell recordings in Fig.1 and 2. Whole cell recordings are from cells near the surface of the slice while field recording sample responses from cells in the middle of the slice.

## Discussion

A new form of synaptic plasticity referred to as presynaptic homeostatic plasticity (PHP) has been described at synapses in the CA1 region of the hippocampus (Chipman *et al*., 2022; Chipman *et al*., 2025). PHP is triggered by the application of the AMPAR antagonist GYKI. It is reported that within minutes of the application of GYKI the depressed AMPAR responses trigger an NMDAR-dependent retrograde presynaptic increase in glutamate release resulting in the recovery of the AMPAR EPSC back to baseline levels. Thus, the two key findings are 1) a recovery of the depressed AMPAR EPSCs in the presence of GYKI and 2) the enhancement of NMDAR EPSCs, interpreted as an increase in glutamate release. Our study uses well established whole cell and field potential recording experiments to address these two findings. We failed to see any recovery of AMPAR responses in the presence GYKI. Second, we saw no increase in the NMDAR EPSC following the application of GYKI.

Thus, we could find no evidence in support of PHP in CA1 hippocampal neurons triggered by GYKI. The present results are seemingly in conflict with the results of Chipman et al. (Chipman *et al*., 2022; Chipman *et al*., 2025). We have gone to great lengths to design the most internally controlled whole cell and field potential experiments, although some differences from the previous studies should be noted. During Chipman et al. experiments they intermittently switched to current-clamp mode. No reason is given for such an unorthodox procedure. Nevertheless, this potential issue does not apply to the results in Fig. 2 and Fig. 3. We used a different pipette solution. However, this would not impact the results in Fig. 2 and 3. Most of our experiments were conducted at room temperature, whereas Chipman et al., used 35 C. However, we repeated the results in Fig. 2 at 35 C and found the same result. Finally, the slicing protocol differed. After 40 years of experience in hippocampal slice physiology, we have settled on the slice protocol used in the present study, as the one that produces the healthiest slices. Nevertheless, we are unaware that the slicing protocol could determine the expression of such a dramatic synaptic phenomenon. Most importantly, as reviewed in the Introduction, 11 studies in which AMPAR transmission was reduced by a wide variety of manipulations and recorded under various conditions none of these studies reported any enhancement in the NMDAR EPSC, an essential requirement for PHP. In summary, based on the present results and the extensive previous literature, we question the existence of PHP at hippocampal CA1 synapses.

## Methods

### Acute Slice Preparation

Acute hippocampal slices were prepared from P65 to P70 mice. Mice were anesthetized with isoflurane. Brains were removed and sliced into 300 μm near-horizontal sections using a Leica VT1200 vibrating microtome. Slices were then transferred to a holding chamber containing ACSF (in mM) (125 NaCl, 2.5 KCl, 1.25 NaH_2_PO_4_, 25 NaHCO_3_, 11 glucose, 1 MgSO_4_, 2 CaCl_2_ saturated with 95% O_2_/5% CO_2_) and incubated for 10 min at 37 ^°^C and then kept at room temperature until use.

### Electrophysiological Recording

All electrophysiological recordings were carried out on an upright Olympus BX51WI microscope and collected using a Multiclamp 700B amplifier (Molecular Devices). During recording, slices were maintained in ACSF (in mM): 125 NaCl, 2.5 KCl, 1.25 NaH_2_PO_4_, 25 NaHCO_3_, 11 glucose saturated with 95% O_2_; 5% CO_2_). Whole cell recording glass patch electrodes filled with an intracellular solution (in mM): 135 CsMeSO_3_, 10 HEPES, 8 NaCl, 0.3 EGTA, 4 Mg-ATP, 0.3 Na-GTP, 5 QX-314, and 0.1 spermine. Excitatory synapses in *Stratum Oriens* were stimulated with a bipolar electrode (Micro Probes) were stimulated with a bipolar electrode (Micro Probes). AMPAR EPSCs were recorded at -70 mV and measured at the peak of the response. NMDAR EPSCs were obtained at +40 mV and measured at 100 ms. Series resistances typically ranged from 10 to 20 MΩ. Most experiments were carried out at 20 to 25 °C. However, an additional series of experiments shown in Fig. 2 were also carried out at 32-34 C. No difference in the results was seen. For field recordings, the internal solution was 3 M NaCl with a large opening pipette tip. The bipolar stimulating electrode (Micro Probes) and recording electrode were placed in either *Stratum Radiatum* or *Stratum Oriens*. AMPAR-mediated responses were collected in the presence of 100 μM picrotoxin to block inhibition. The slope of the field EPSPs was used to measure response size. Statistical difference was determined using a two-tailed paired t test.

